# Normobaric hypoxia alters the transcriptional response of healthy human skeletal muscles to a single session of high-intensity interval exercise

**DOI:** 10.64898/2026.04.22.720051

**Authors:** Jia Li, Dale F. Taylor, Jujiao Kuang, Zhenhuan Wang, Navabeh Zare, Muhammed M. Atakan, Kangli Cui, Nima Ouzhu, Bianba, Andrew Garnham, Wentao Lin, Li Peng, Olivier Girard, David J. Bishop, Yanchun Li, Xu Yan

**Affiliations:** College of Physical Education, Southwest University, Chongqing, China; Institute for Health and Sport, Victoria University, Melbourne, Australia; Australia Institute for Musculoskeletal Sciences, Melbourne, Australia; Division of Exercise Nutrition and Metabolism, Faculty of Sport Sciences, Hacettepe University, Ankara, Türkiye; College of Education, Tibet University, Lhasa, Tibet, China; High Altitude Medical Research Center, Medical College, Tibet University, Lhasa, Tibet, China; College of Sport Science, Zhuhai College of Science and Technology; School of Human Sciences (Exercise and Sport Science), The University of Western Australia, Perth, Australia; China Institute of Sport and Health Science, Beijing Sport University, Beijing, China; Department of Medicine-Western Health, The University of Melbourne, Melbourne, Australia

## Abstract

Given its well-documented effects on human physiology, hypoxia has garnered increasing interest for its potential to enhance specific adaptations to exercise. However, the molecular response of skeletal muscle to exercise under normobaric hypoxia remains poorly understood. To address this gap in knowledge, ten healthy young males completed a crossover study in which exercise in hypoxia was compared to exercise in normoxia matched by either absolute or relative intensity. This design allowed us to identify shared transcriptomic responses across all three conditions, as well as changes that were specific to exercise intensity or hypoxic exposure. Skeletal muscle biopsies were collected before, immediately after, and at 3 and 24 hours following each exercise session, with RNA sequencing performed to assess changes in gene expression. Following exercise, a greater number of differentially expressed genes were observed in hypoxia compared to normoxia at 24 h post-exercise. This hypoxia-specific response involved the downregulation of multiple mitochondrial pathways and appears to be regulated by a transcriptional network comprising both positive and negative regulators of HIF-1 activity. These findings highlight the ability of normobaric hypoxia to influence exercise-induced gene expression and suggests that it may promote distinct molecular adaptations in skeletal muscle following longer-term training.

## Introduction

Exercise is a potent non-pharmacological intervention that induces numerous health benefits^1^. By adjusting variables such as exercise intensity, duration, and modality, specific molecular and physiological adaptations can be induced^2^. To amplify training-induced responses, there is increasing interest in modifying environmental factors^3^. For example, hypoxia exposure, characterised by decreased oxygen availability and supply to tissues^4^, has long been associated with greater improvements in performance through increased red blood cell production^5^, greater capillary density^6^, and changes in mitochondrial respiratory function^7,8^. However, the potential benefits of intermittent hypoxic training, where exercise occurs in hypoxia but without chronic exposure, remain poorly understood.

Exercising in hypoxia is thought to increase metabolic stress compared to normoxia, potentially enhancing training-induced adaptations from cardiorespiratory exercise. However, comparing exercise performed with and without hypoxia is challenging because common exercise prescription metrics, such as peak power output (PPO)^9^ and peak rate of oxygen consumption attainable during physical exertion (V̇O_2peak_)^10^, are lower in hypoxia due to reduced oxygen availability. To standardise the training stimulus, exercise in normoxia is typically matched to either the absolute or relative intensity used in hypoxia^11^. These differing approaches have led to inconsistent findings regarding the potential benefit of exercising in hypoxia. Studies generally report greater improvements or no difference in fitness gains depending on whether exercise in normoxia was matched by absolute or relative intensity, respectively^12^. However, the limited number of studies, small sample sizes, differences in hypoxic exposure, and inconsistencies in exercise matching make it difficult to draw firm conclusions. Nonetheless, hypoxia has been shown to influence the expression of genes involved in energy metabolism, both in response to a single session of exercise^13^ and following training^14^, as well as induce^15^ or enhance^16,17^ specific mitochondrial adaptations. As such, changes in skeletal muscle physiology may still occur from hypoxia even when improvements in whole-body metrics, such as V̇O_2peak_, are not evident.

The remarkable plasticity of skeletal muscle underscores improvements in exercise capacity and metabolic health, with cellular modifications varying according to the stimulus provided^18^. In hypoxia, the cellular response is primarily regulated by hypoxia-inducible factor 1α (HIF-1α) – a transcription factor that is rapidly degraded in normoxia but stabilised in low-oxygen conditions^19^. Both exercise^13,20^ and hypoxic exposure^21^ increase HIF-1α protein abundance in skeletal muscle, facilitating gene expression related to angiogenesis and energy metabolism through binding to hypoxia-inducible factor 1β (HIF-1β, often called ARNT) to form HIF-1. In contrast, HIF-1 has an inhibitory effect on transcription factors such as MYC^22^ and peroxisome proliferator-activated receptor gamma coactivator 1-alpha (PGC-1α)^23^, both of which regulate mitochondrial gene expression. Despite the complex effects of hypoxia on the regulation of gene expression^22^, no study has investigated how hypoxia influences global exercise-induced gene expression in skeletal muscle following a single exercise session using RNA sequencing (RNA-seq).

To fill this knowledge gap, this study aimed to investigate the transcriptional response to a single session of high-intensity interval exercise (HIIE) in hypoxia, with exercise matched to absolute or relative intensity in normoxic conditions. Using a crossover study design, participants completed three HIIE sessions, with four muscle biopsies collected before (PRE), immediately after (+0 h), and at 3 (+3 h) and 24 hours (+24 h) following exercise. This powerful approach provided a comprehensive assessment of the combined effects of normobaric hypoxia and HIIE on human skeletal muscle, allowing us to investigate a shared exercise response across all three conditions, as well as intensity-specific and hypoxia-specific changes in the transcriptome. By elucidating how hypoxia alters the transcriptional response to exercise, more precise strategies may be developed for enhancing athletic performance and preventing or treating exercise-responsive diseases.

## Results

### Hypoxia induces a distinct transcriptional response to both absolute and relative intensity-matched normoxic exercise

Although hypoxia is known to modulate exercise-induced gene expression in skeletal muscle, previous assessments have relied on targeted analyses using quantitative PCR (qPCR)^13,21,24,25^. Additionally, most studies have not included comparisons to both absolute and relative intensity-matched exercise performed in normoxia^12^, thereby limiting the ability to disentangle adaptations that are specific to hypoxia and those that are induced by differences in intensity between exercise prescriptions. To address this, RNA-seq was performed on skeletal muscle biopsies collected from 10 healthy young males before, immediately after, and 3 and 24 hours following HIIE in hypoxia (HY) and normoxic sessions matched for either the same absolute power (NA) or the same relative intensity (NR) as HY (**Fig. 1a-b**). This protocol resulted in a significant difference in power between NA and HY compared to NR (**Supplementary Fig. 1a**), as well as in the percentage of PPO (**Supplementary Fig. 1b**) and the percentage of maximal heart rate achieved (**Fig. 1c**) for HY and NR compared to NA, but not between HY and NR. Successful matching of both absolute and relative exercise intensity between normoxic and hypoxic conditions was confirmed by comparable post-exercise blood lactate elevations in HY and NR, both of which exceeded those observed in NA (**Fig. 1d**). However, only HY elicited a significant post-exercise rise in blood glucose, indicating a unique metabolic stress induced by hypoxia (**Fig. 1e**).

**Fig. 1.**
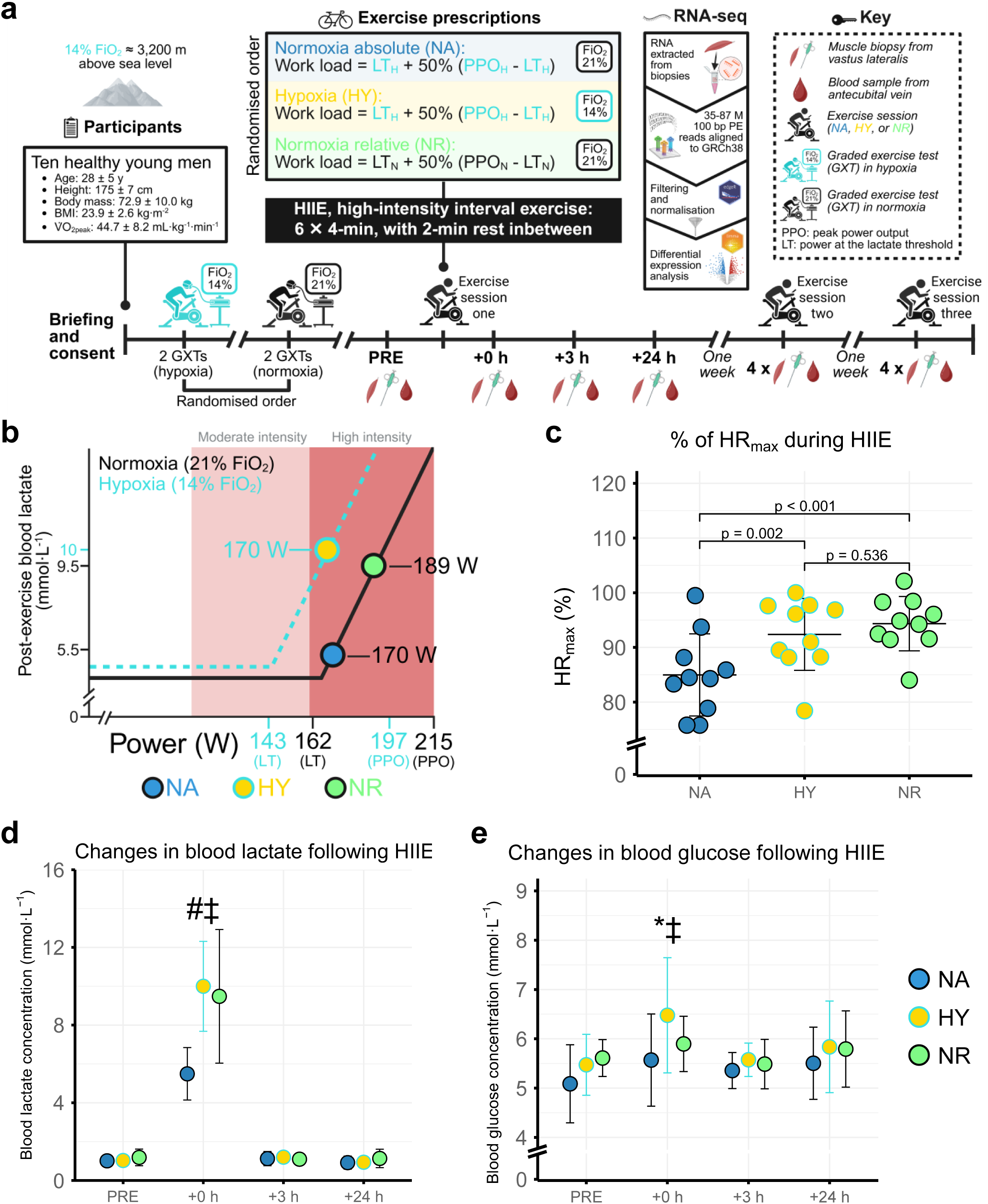
Study overview. **a** Study design, including baseline exercise testing using graded exercise tests (GXTs) conducted in both normoxia and hypoxia, prescription of a single session of high-intensity interval exercise (HIIE) in hypoxia (HY) and normoxia, matched by absolute (NA) and relative intensity (NR), muscle biopsy and blood collection, and bioinformatics workflow. PE: paired-end. **b** Prescription of high-intensity interval exercise for NA, HY, and NR, relative to peak power output (PPO) and the lactate threshold (LT) in normoxia and hypoxia. Values displayed are group averages. **c** Differences in the % of heart rate max (HR_max_) achieved between NA, NR, and HY, with HR_max_ identified from either the GXT in normoxia or hypoxia. **d** Changes in antecubital blood lactate following HIIE in NA, HY, and NR. All conditions had blood lactate significantly increase from PRE to +0 h, however, a significant difference was observed between the blood lactate at +0 h in NR and HY to NA (indicated by # and ‡ respectively). **e** Changes in antecubital blood glucose following HIIE in NA, HY, and NR. A significant increase was observed from PRE to +0 h only in HY (indicated by *) and a significant difference between the blood glucose at +0 h in HY to NA (indicated by ‡). *, #, and ‡ indicates p < 0.05.

Skeletal muscle biopsies were obtained pre-exercise, immediately post-exercise, and 3 and 24 hours after each HIIE session, after which RNA was extracted and sequenced. Following data pre-processing and normalisation, 16,352 transcripts remained for quantification. This included 96% (12,466 of 12,945) of protein-encoding genes annotated in the Human Protein Atlas^26^ (**Supplementary Fig. 2a**). Multidimensional scaling (MDS) revealed clustering of the PRE and +0 h time points together, with distinct clusters for +3 h and +24 h also observed (**Fig. 2a**). As expected, no genes were differentially expressed between the PRE time points across the three conditions, confirming a return to baseline gene expression between exercise trials. Consistent with the MDS plot and previous studies using HIIE^27^, few genes were differentially expressed at +0 h relative to PRE in any condition (**Fig. 2b**). Reflecting the influence of relative exercise intensity on early gene expression^28^, the NA condition showed the fewest number of differentially expressed genes (DEGs) at +3 h relative to PRE, whereas the HY and NR conditions exhibited a similar number of upregulated DEGs. However, despite matching for relative intensity, the HY condition displayed a greater number of DEGs at +24 h (8,250 DEGs) compared to NR (4,827 DEGs), indicating a delayed, hypoxia-specific gene expression response to HIIE. When comparing genes upregulated or downregulated at each time point relative to PRE across NA, HY, and NR, the four largest groups were: upregulated in all conditions at +24 h, downregulated only in HY at +24 h, downregulated in all conditions at +24 h, and upregulated only in HY at +24 h (**Fig. 2c**).

**Fig. 2.**
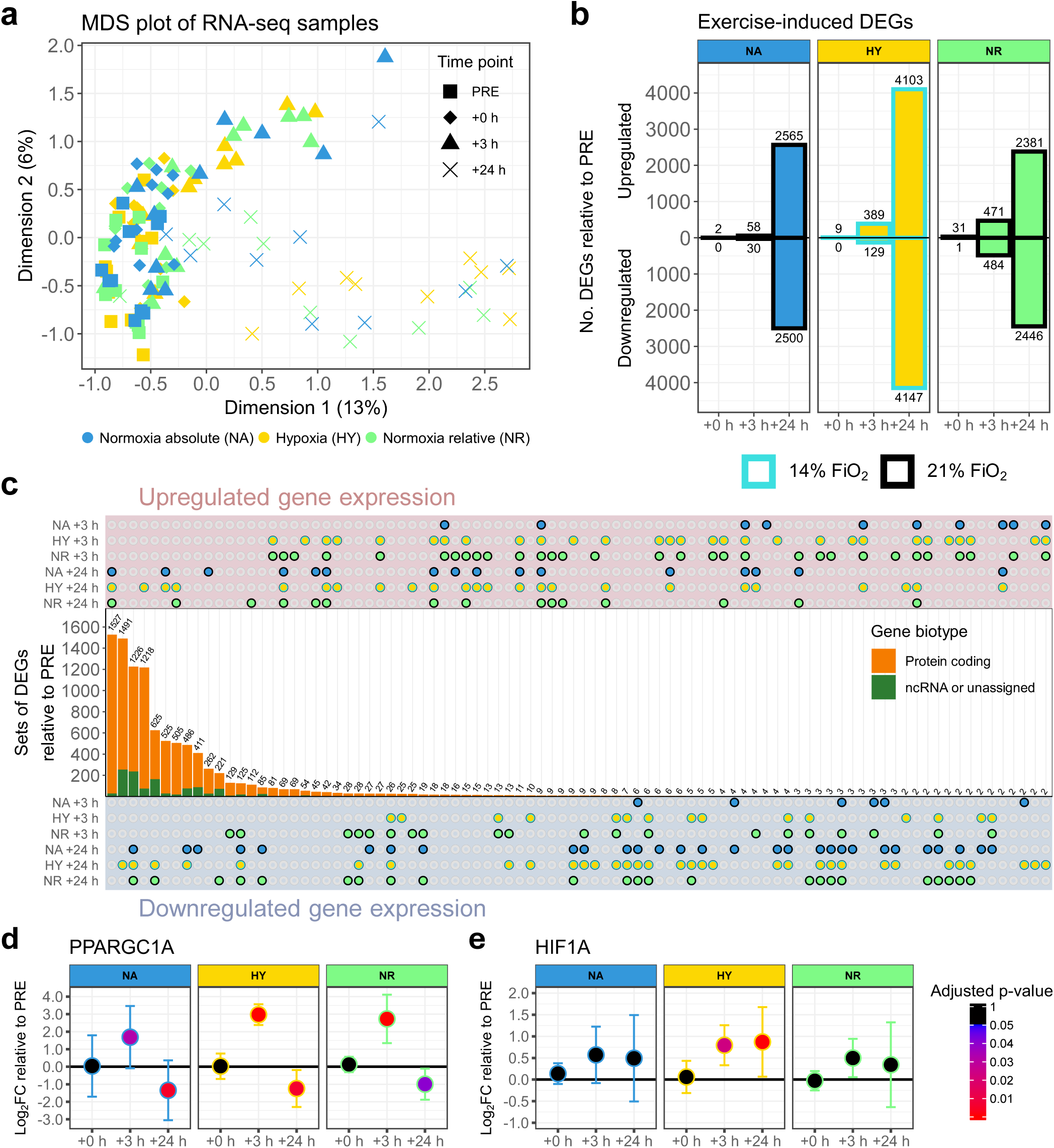
Effect of hypoxia on exercise-induced gene expression compared to absolute and intensity-matched exercise in normoxia. **a** Multidimensional scaling (MDS) of CPM values following data filtering and normalisation. **b** The number of upregulated and downregulated differentially expressed genes (DEGs) relative to PRE at each time point in NA, HY, and NR. Significance was determined with an adjusted p-value < 0.05 (Benjamini-Hochberg (BH)). **c** Upset plot of upregulated and downregulated DEGs at each time point relative to PRE in NA, HY, and NR, with the proportion of each set that is protein-coding indicated. Minimum set size displayed is 2. ncRNA: non-coding RNA. **d** Differential expression relative to PRE of *PPARGC1A*, with upregulation at +3 h and downregulation at +24 h in NA, HY, and NR. Significance was determined with an adjusted p-value < 0.05 (BH). **e** Differential expression relative to PRE of *HIF1A*, with upregulation at +3 h and +24 h only in HY. Significance was determined with an adjusted p-value < 0.05 (BH).

Differentially expressed genes shared across all conditions included the commonly studied regulator of mitochondrial biogenesis, *PPARGC1A* (encoding PGC-1α). *PPARGC1A* was upregulated at +3 h in all conditions, but the magnitude of upregulation was lower in NA, consistent with a lower relative exercise intensity^28^ (**Fig. 2d**). In contrast, *HIF1A* was significantly upregulated exclusively in HY at +3 h and +24 h relative to PRE (**Fig. 2e**), consistent with previous reports of its enhanced transcription under hypoxic conditions^13,20^. Together, these results suggest that specific genes or pathways may be regulated in an intensity- or hypoxia-specific manner.

### Exercise induces pathway-specific differential expression in an intensity- and hypoxia-dependent manner

While it is widely accepted that the transcriptional response to exercise is influenced by the various parameters used to prescribe exercise, such as intensity^28–31^, it is also likely that a core exercise-induced transcriptional response exists that is largely independent of the specific exercise prescription^30,32^. We leveraged our experimental model to determine whether a shared transcriptional response was common across all exercise conditions. To achieve this, enrichment analysis using the gene ontology biological process (GOBP) database was performed for upregulated and downregulated DEGs at each time point shared between NA, HY, and NR (**Fig. 3a**). As expected, no terms were enriched at +0 h (1 shared DEG) or +3 h (44 shared DEGs), largely due to the low number of DEGs in the NA condition. At +24 h, a substantial number of enriched terms were observed (3,078 shared DEGs), especially among upregulated genes, including those associated with DNA replication, methylation, ribosome biogenesis, nuclear export, protein folding, autophagy, and vesicle organisation. In contrast, the shared downregulated terms were predominantly linked to skeletal muscle development, such as extracellular matrix organisation and muscle organ development. However, even among genes that were differentially expressed across all conditions, the distribution of log₂ fold changes was most strongly shifted away from zero in HY, followed by NR and then NA at +3 h. By +24 h, when a substantially larger set of shared DEGs was detected, this shift was significantly greater in HY compared with both NA and NR, for both upregulated and downregulated genes (**Fig. 3b**). These findings suggest the presence of hypoxia-specific mechanisms that amplify the transcriptomic response to a single session of HIIE.

**Fig. 3.**
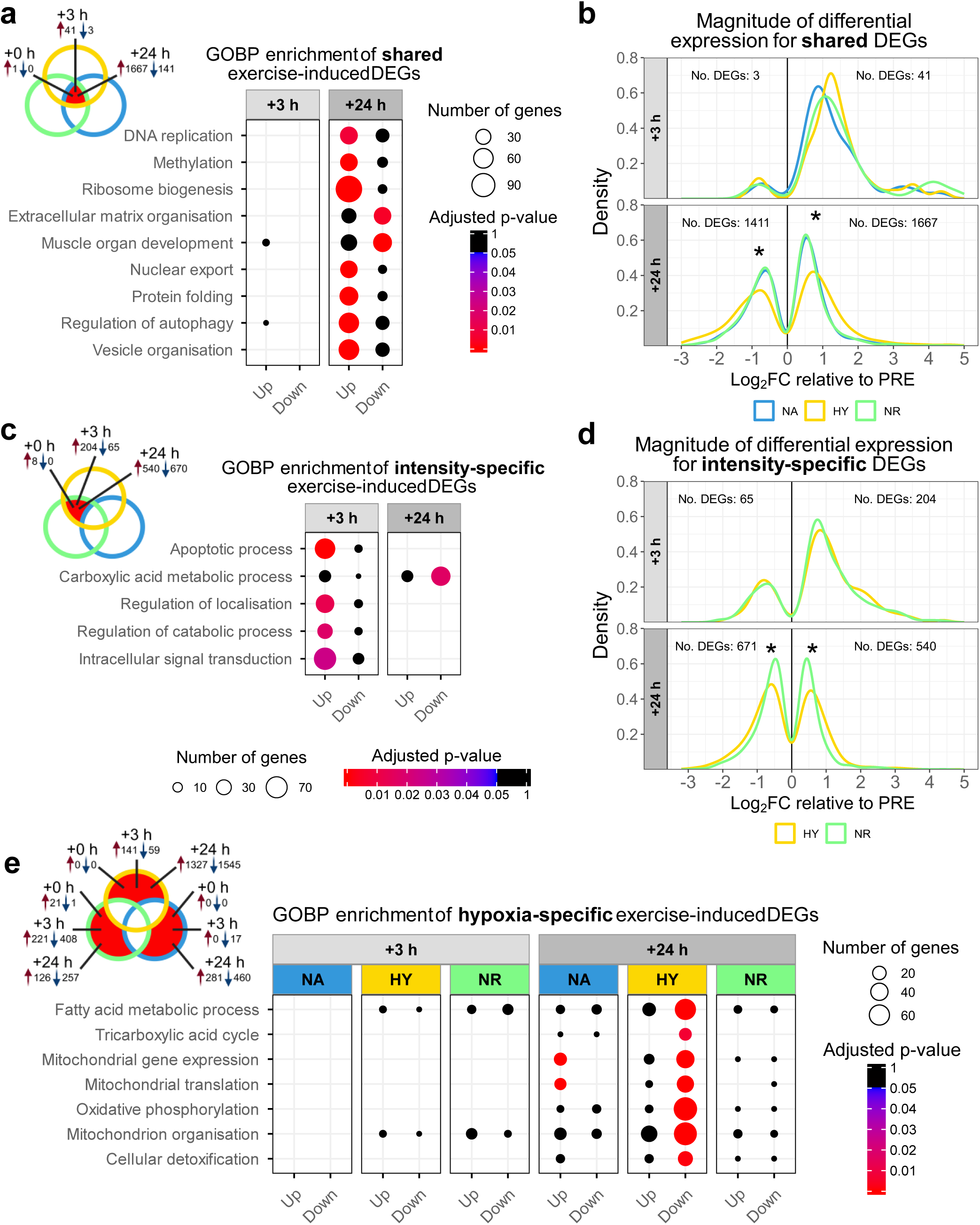
Shared, intensity-specific, and hypoxia-specific pathways induced by exercise. **a** Gene ontology biological process (GOBP) enrichment of differentially expressed genes (DEGs) with shared upregulation or downregulation at each time point relative to PRE between NA, HY, and NR. Significant enrichment was determined with an adjusted p-value < 0.05 (Benjamini-Hochberg (BH)). **b** Ridge plot illustrating the log_2_ fold change (FC) densities of DEGs shared between all conditions at each time point relative to PRE. A significant difference (adjusted p-value < 0.05 (Holm-Bonferroni)) was observed between HY and NA, and between HY and NR, for both downregulated and upregulated genes at +24 h (indicated by *). **c** GOBP enrichment of DEGs with shared upregulation or downregulation at each time point relative to PRE between HY and NR, but not NA. Significant enrichment was determined with an adjusted p-value < 0.05 (BH). **d** Ridge plot illustrating the log_2_ FC densities of DEGs shared between HY and NR, but not NA at each time point relative to PRE. A significant difference (adjusted p-value < 0.05 (Holm-Bonferroni)) was observed between HY and NA, and between HY and NR, for both downregulated and upregulated genes at +24 h (indicated by *). **e** GOBP enrichment of DEGs at each time point relative to PRE uniquely upregulated or downregulated in NA, HY, and NR. Only enrichments that were significant in HY are displayed. Significant enrichment was determined with an adjusted p-value < 0.05 (BH).

To investigate intensity-dependent responses to exercise, enrichment analysis using the GOBP database was performed for upregulated and downregulated DEGs at each time point shared between HY and NR, but not NA (**Fig. 3c**). No pathways were shared at +0 h, consistent with the limited number of DEGs observed across the two conditions. At +3 h, enrichment of upregulated apoptotic processes, alongside broader terms related to signal transduction, the regulation of localisation, and the regulation of catabolism, were observed between HY and NR, suggesting regulation in an intensity-specific manner. At +24 h, several terms associated with organic acid metabolism were identified as downregulated between HY and NR. Among genes shared between only HY and NR, the distribution of log₂ fold changes was most strongly shifted away from zero in HY once again, with a significant difference observed for both upregulated and downregulated genes at +24 h (**Fig. 3d**), further indicating hypoxia-specific effects on exercise-induced gene expression.

As both the shared exercise response and the intensity-specific response indicate a moderating effect of hypoxia, we next investigated the hypoxia-specific response to exercise. To do this, enrichment analysis using the GOBP database was performed for upregulated and downregulated DEGs at each time point that were unique to hypoxia, as well as those unique to NA and NR – representing exercise-responsive genes whose differential expression was absent under hypoxic conditions (**Fig. 3e**). As few genes were shared exclusively between NA and NR, these were excluded from the visualisation. As expected, no terms were enriched at +0 h post-exercise, given the minimal number of DEGs across all conditions. The largest number of significant terms was observed for HY at +24 h, with many related to the mitochondrion identified, including mitochondrial gene expression, translation, organisation, and several related to energy generation, such as oxidative phosphorylation (OXPHOS). The mitochondrial enrichments in HY were for downregulated genes and contrasted to an enrichment of upregulated genes involved in mitochondrial gene expression and translation in NA. As such, these results indicate a potential combined intensity- and hypoxia-specific modulation of exercise-induced mitochondrial gene expression.

### Hypoxia strongly amplifies the downregulation of nuclear-encoded genes involved in oxidative phosphorylation

To better understand the intensity- and hypoxia-specific effect on different mitochondrial pathways, the distribution of log_2_ fold changes for some of the significantly enriched terms from the GOBP enrichment analysis were visualised. Consistent with the GOBP enrichment results, the distributions of log₂ fold changes for genes associated with mitochondrial gene expression (**Fig. 4a**), mitochondrial translation (**Fig. 4b**), and mitochondrial organisation (**Fig. 4c**) were shifted toward more negative values in HY compared with both NA and NR, which showed similar patterns in both the number of differentially expressed genes and the magnitude of change. Genes involved in oxidative phosphorylation were also distinctly shifted leftward relative to NR at +3 h and +24 h, and even more so relative to NA at +24 h (**Fig. 4d**), albeit with no substantial change in the number of DEGs between the latter two conditions.

**Fig. 4.**
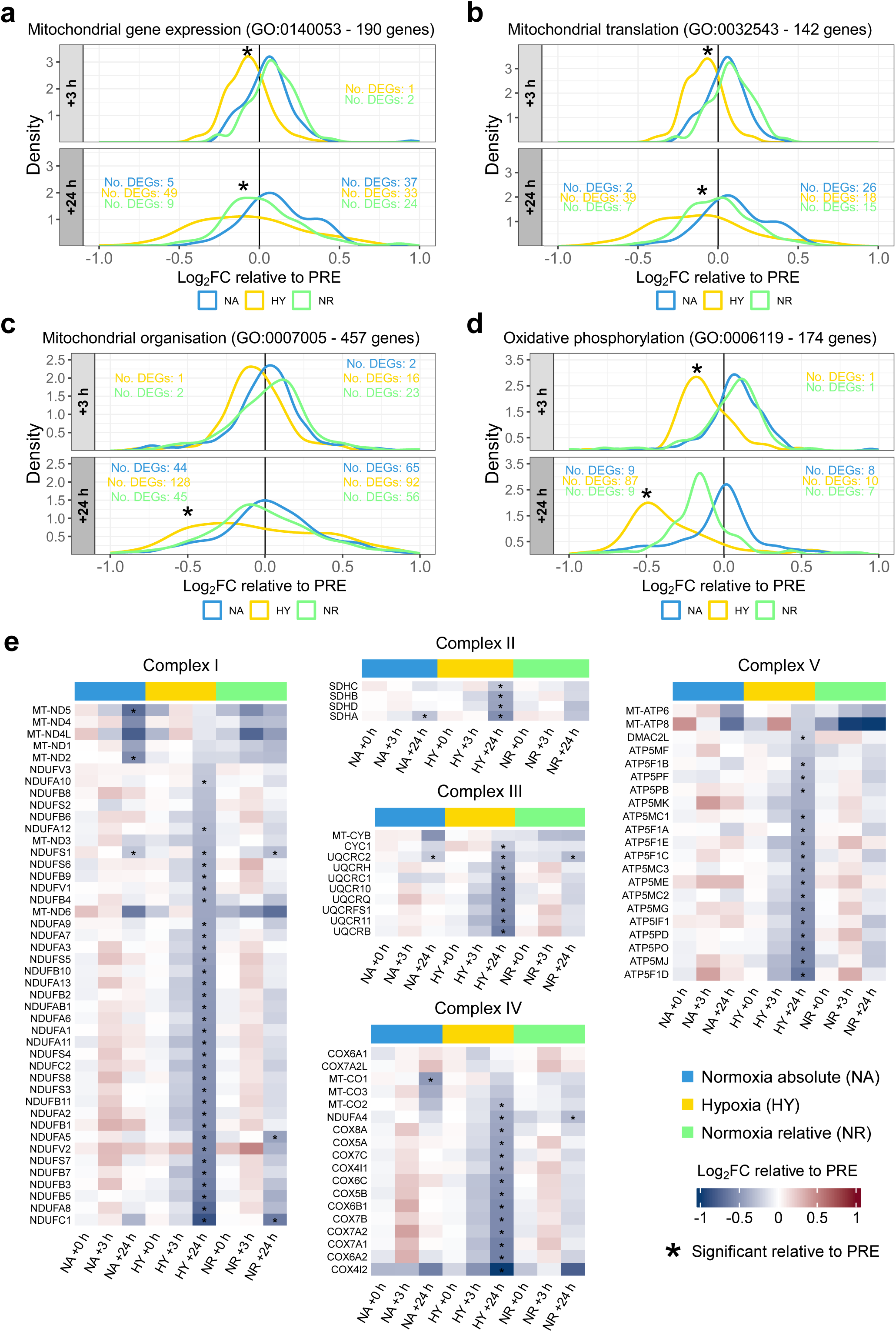
Hypoxic-specific downregulation of mitochondrial genes in response to exercise. Ridge plots illustrating the log_2_ fold change (FC) densities at each time point relative to PRE for gene ontology biological process (GOBP) terms: **a** Mitochondrial gene expression. **b** Mitochondrial translation. **c** Mitochondrial organisation. **d** Oxidative phosphorylation. A significant difference (adjusted p-value < 0.05 (Holm-Bonferroni)) between HY and NA, and between HY and NR, for all quantified genes is indicated by *. **e** Heatmap of genes in oxidative phosphorylation complexes I, II, III, IV, and V. Significance was determined with an adjusted p-value < 0.05 (Benjamini-Hochberg) and is indicated by * for each combination of condition and time point.

Examination of individual genes involved in oxidative phosphorylation revealed that 77 of the 87 downregulated genes at +24 h in HY encoded subunits of the five complexes of the electron transport chain (**Fig. 4e**). Few of these genes reached statistical significance at other time points in HY, or in either NA or NR at any time point relative to PRE. An exception were some mitochondrial DNA-encoded subunits, which exhibited downregulation only in NA at +24 h, but were largely unaffected in HY. Collectively, these results suggest that hypoxia primarily modulates the delayed transcriptional response of nuclear-encoded mitochondrial genes to exercise, potentially mediated by altering the activation of specific nuclear transcription factors.

### The hypoxia-specific exercise response likely involves upregulation of both activators and repressors of HIF-1 activity

To explore potential transcriptional regulators underlying the transcriptomic response to exercise in hypoxia, transcription factor activity was inferred using DoRothEA regulons with the VIPER method^33^. This is a well-established method that uses filtered and normalised gene expression data to infer changes in transcription factor activity by integrating both the direction and magnitude of expression changes across high-confidence transcription factor–target gene regulons. Reflecting the amplified transcriptomic response at +24 h, more transcription factors were predicted to have upregulated activity at +24 h relative to PRE in hypoxia compared to the two exercise conditions in normoxia (**Fig. 5a**). Unlike gene expression, HIF-1α transcriptional activity was predicted to be upregulated at +3 h across all conditions, consistent with previous research investigating the response to a single session of exercise^20^ (**Fig 5b**).

**Fig. 5.**
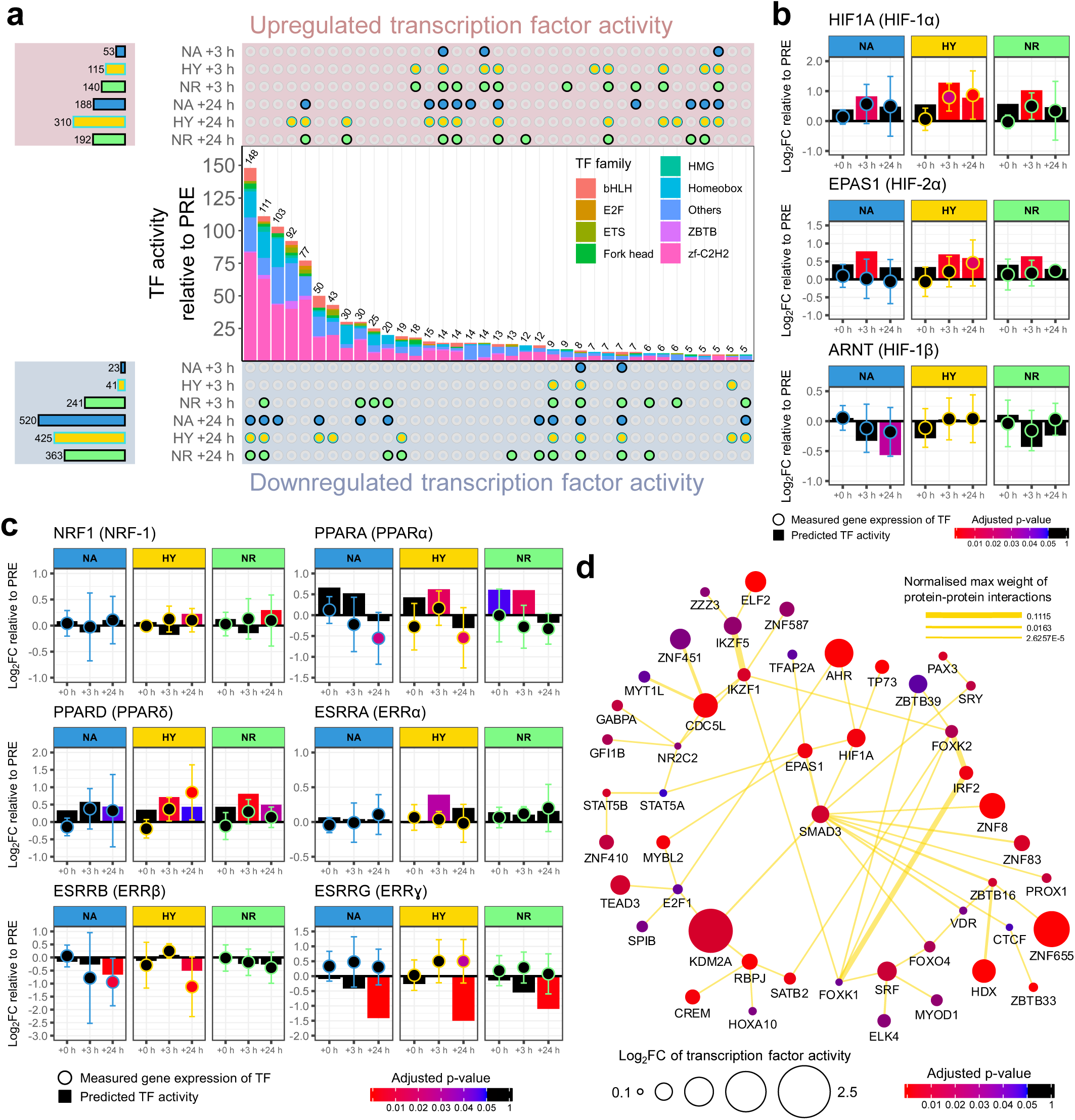
Identification of transcription factors activated by exercise in hypoxia. **a** Upset plot of transcription factors (TF) with differential activity relative to PRE identified using DoRothEa regulons and the VIPER method^33^. Significance was determined with an adjusted p-value < 0.05 (Benjamini-Hochberg). Minimum set size displayed is 5. bHLH: basic helix-loop-helix. HMG: high mobility group. ZBTB: Zinc finger and BTB domain containing. Zf-C2H2: Cys2–His2 zinc finger. **b** Differential gene expression and transcription factor activity of HIF-1α and EPAS1 (HIF-2α), and their binding partner ARNT (HIF-1β). **c** Differential gene expression and transcription factor activity of transcription factors known to regulate mitochondrial gene expression. **c** Protein-protein physical interaction network using GeneMANIA^36^ for transcription factors with activity upregulated within only HY at +24 h using DoRothEa regulons and the VIPER method. A minimum interaction confidence of 0.4 was required for visualisation and all transcription factors without a physical interaction to another enriched transcription factor were excluded from the visualisation. FC: fold change.

At +24 h, HIF-1α and EPAS1 (HIF-2α), another oxygen-sensing transcription factor, remained upregulated exclusively in HY, suggesting prolonged activation following exercise, even after returning to normoxic conditions following exercise completion. In contrast, the activity of their binding partner ARNT (HIF-1β), which is required for HIF-1 and HIF-2 to induce transcription, remained unchanged following exercise in hypoxia.

Despite the downregulation of mitochondrial genes following exercise in hypoxia, transcription factors commonly linked to mitochondrial biogenesis^34^ – the synthesis of new components of the mitochondrial reticulum, showed similar predicted activity across all conditions (**Fig. 5c**). Aside from the potential involvement of other regulatory mechanisms, such as epigenetic modifications or altered chromatin accessibility^35^, this indicates that mitochondrial gene expression may be downregulated through separate transcription factors. To uncover the transcriptional network driving the hypoxia-specific response, protein-protein interactions among transcription factors with predicted upregulated activity at +24 h in hypoxia were mapped using GeneMANIA^36^ (**Fig. 5d**), with differential activity and the strength of physical associations (via the STRING database) calculated^36^. Many of these transcription factors clustered together as one cohesive network. In addition to HIF-1α and HIF-2α, this network included transcription factors with previously identified roles in regulating the response to hypoxia, including SMAD3^37^, AHR^38^, TFAP2A^39^, TP73^40^, and HDX^41^, among others. Together, these findings highlight a coordinated transcriptional response to exercise in hypoxia, driven by a network of transcription factors that extends beyond the canonical HIF-1 pathway.

## Discussion

Using two physiologically distinct normoxic exercise prescriptions, we demonstrate that exercise in hypoxia induces a unique transcriptional response, independent of both absolute and relative exercise intensity. This hypoxia-specific response, most prominent 24 h post-exercise, was characterised by an amplification of the DEGs shared across all three conditions, along with an amplification of the intensity-specific DEGs altered in only HY and NR. In addition, a distinct set of genes was differentially expressed exclusively under hypoxic conditions. Enrichment analysis of HY-specific DEGs identified mitochondrial pathways as significantly overrepresented, with genes involved in oxidative phosphorylation notably more downregulated at +24 h. To identify the regulatory mechanisms underlying these transcriptional changes, transcription factor activity analysis at this time point revealed a hypoxia-specific regulatory network enriched for both positive and negative modulators of HIF-1 activity. Together, these findings underscore the ability of normobaric hypoxia to modulate exercise-induced gene expression.

Although the absolute and relative exercise intensities and environmental oxygen levels differed between conditions, each elicited the differential expression of thousands of overlapping genes (**Fig. 2c**). This overlapping response may therefore represent a common contraction-driven response to exercise that is largely independent of the specific characteristics of the exercise stimulus, an idea that has previously received only limited investigation^30,32^. It is worth nothing, however, that all three exercise intensities could be classified as high (i.e., between the second metabolic threshold and V̇O_2peak_)^42^, and it is possible that this common contraction-driven transcriptional response may have been overestimated due to the relatively narrow range of exercise intensities. Nonetheless, this shared transcriptomic response, predominantly observed at +24 h and characterised by a higher proportion of upregulated than downregulated genes, aligns with previous comparisons of exercise-induced gene expression across very different exercise intensities, such as moderate-intensity continuous exercise (MICE) and sprint interval exercise (SIE)^31^ (**Supplementary Fig. 3a**). Indeed, 70% of the shared upregulated genes at +24 h (compared with only 42% of shared downregulated genes at +24 h) in our study were also differentially expressed following MICE and SIE, two exercise intensities substantially below and above our conditions (see **Supplementary Fig. 3b** for changes in lactate following exercise and **Supplementary Fig. 3c** for differences in exercise volume between all five conditions), respectively, despite differences in gene coverage. For the shared upregulated genes in our study (**Fig. 3a**), an enrichment of numerous GOBP terms was observed, including those related to DNA replication, methylation, ribosome biogenesis, nuclear export, protein folding, regulation of autophagy, and vesicle organisation. Further supporting these pathways as part of a common contraction-driven response to exercise, nearly all of the same pathways, with the exception of DNA replication and extracellular matrix organisation, remained enriched when genes differentially expressed following MICE and SIE were also incorporated (**Supplementary Fig. 3d**). These pathways are likely required for adaptation regardless of exercise modality, with protein translation via ribosomal biogenesis and protein folding required for processes driven by distinct and sometimes opposing signalling pathways, such as mitochondrial biogenesis and myofibrillar synthesis^43^. Similarly, autophagy-mediated recycling of damaged cellular components has been observed not only across different intensities of cardiorespiratory exercise, but also between cardiorespiratory and resistance exercise^44^. For the GOBP terms enriched among shared downregulated genes (**Fig. 3a**), the suppression of those related to extracellular matrix organisation and muscle organ development may initially appear unexpected and counterproductive^45^. However, the timing of the muscle biopsy likely does not capture the full temporal dynamics of these genes’ responses to exercise^27,46^. Notably, one study reported transient upregulation of many of these genes at earlier time points following a similar HIIE protocol in normoxia^27^. Additional pathways may also contribute to this shared contraction-driven response but were not detected in the present study because they were not differentially expressed at +3 h or +24 h. Collectively, these findings support the existence of a conserved, contraction-driven transcriptomic response that may be largely independent of exercise prescription. Identifying this shared response is important because it helps distinguish universal molecular adaptations from those specific to a particular exercise intensity, modality, or environmental condition.

For intensity-specific upregulated genes, a smaller number of enrichments was observed between only HY and NR than shared between all conditions (**Fig. 3c**). The limited number of intensity-specific terms, relative to shared terms, may reflect the comparatively small differences in exercise intensity between NA, NR, and HY, as all protocols could be categorised as high-intensity (i.e., between the 2^nd^ metabolic threshold and V̇O_2peak_)^42^. Of the terms identified, a greater upregulation of apoptosis with greater intensity is consistent with observations in lymphocytes^47,48^, with a shift toward anti-apoptotic adaptation generally observed in skeletal muscle following cardiorespiratory training^49^. The downregulation of genes related to carboxylic acid metabolism is consistent with a greater reliance on carbohydrate metabolism at higher exercise intensities and associated changes at the metabolite level^50^, potentially reflecting a negative feedback loop at the gene level. Similarly, these changes are likely further amplified by hypoxia, with intensity-specific upregulated and downregulated genes exhibiting greater log₂ fold changes in hypoxia at +24 h (**Fig. 3d**), mirroring the pattern seen in genes shared across all conditions (**Fig. 3b**).

Following HIIE, hypoxia elicited a greater number of differentially expressed genes compared to exercise matched for either absolute or relative intensity in normoxia (**Fig. 2b**). The additive effect of hypoxia and exercise, both in terms of the number of DEGs and the magnitude of differential expression, is consistent with previous qPCR findings^13,21^. Among these changes, mitochondrial gene expression was notably downregulated, suggesting a potential impact on subsequent mitochondrial adaptations. This result contrasts with some previous studies on exercise training in normobaric hypoxia. These studies generally report additional effects following a period of training on measures such as mitochondrial volume density^14,16,51,52^ and citrate synthase (CS) activity^14,16,51,52^ when exercise in normoxia is matched by absolute intensity. However, when matched by relative intensity, studies have generally reported no differences for changes in electron transport chain subunit protein abundance^53^, mitochondrial respiratory function^54^, or CS activity^6,7,55^, or even a negative effect on changes in CS activity^56^ and mitochondrial respiratory function^57^, although some increases in mitochondrial volume density have been observed^14,52^. Greater improvements in mitochondrial volume density with hypoxic compared to normoxic exercise matched by relative intensity were observed only in high-intensity exercise groups (65.6% vs. 67% of maximum power output (Ẇ_max_) respectively^14^ or 85% of heart rate max (HR_max_)^52^), but not in moderate-intensity groups (52.4% vs. 57.8% of Ẇ_max_ respectively or 77% of HR_max_), for each study. Thus, an augmentation of mitochondrial adaptations from hypoxic exercise may be intensity-specific and affect only certain mitochondrial characteristics. Although one might speculate that the downregulation of mitochondrial genes in HY indicates a blunting of mitochondrial adaptations under our exercise prescription, the concurrent upregulation of several of the same pathways in NA complicates this interpretation (**Fig. 3e**). Indeed, higher exercise intensities and volume are generally associated with greater increases in mitochondrial content and respiratory function^2^. These results may instead reflect a ‘first-bout’ effect, in which downregulation of HIF-1 signalling, potentially via increased expression of its negative regulators, represents an early adaptation that leads to altered exercise-induced gene expression in subsequent sessions^58,59^.

While the role of HIF-1 in regulating hypoxic responses is well established^60^, the contribution of other transcription factors and co-activators, particularly when combined with exercise, remains poorly characterised. Our results indicate that hypoxia activates multiple transcription factors that act to either upregulate or downregulate pathways induced by HIF-1. Consistent with previous research, SMAD3 emerged as a central hub in the hypoxic response^37^, physically interacting with other enriched transcription factors, including HIF-1α and HIF-2α (**Fig. 5d**). As a downstream effector of TGF-β cytokines, SMAD3 has been implicated in repressing *PPARGC1A* gene expression in skeletal muscle^61^, which we observed at +24 h across all conditions. PGC-1α is an essential co-activator of many key regulators of mitochondrial gene expression^34^, including members of the nuclear respiratory factor (NRF), peroxisome proliferator-activated receptor (PPAR), and estrogen-related receptor (ERR) transcription factor families^34^. As such, downregulation of PGC-1α activity may contribute to the altered mitochondrial gene expression following hypoxic exercise and support the HIF1-driven shift toward glycolytic metabolism^13,62^ – a pattern reflected in our results, where a larger number of genes encoding glycolytic enzymes were differentially expressed, or showed greater upregulation, in HY (**Supplementary Fig. 4**). In addition to oxygen-sensitive transcription factors, KDM2A was identified as binding to SMAD3 and exhibited the greatest fold change among transcription factors uniquely upregulated in the hypoxia-specific network. KDM2A, a lysine-specific histone demethylase, has been implicated in suppressing *ESRRG* gene expression and the activity of its encoded transcription factor, ERRγ^63^. Additionally, inhibition or deletion of KDM2A has been shown to promote a shift toward greater aerobic metabolism in skeletal muscle, suggesting a potential role for hypoxia in downregulating mitochondrial biogenesis^63^. However, whether this regulation occurs following exercise in hypoxia requires experimental validation. Several transcription factors that negatively regulate HIF-1α activity, such as AHR, which competes with HIF-1α for ARNT binding^38^, and TP73, which facilitates HIF-1α degradation^40^, were also upregulated exclusively under the hypoxia condition. Counterintuitively, transcription factors like ZBTB16^64^ and GABPA (also known as NRF-2)^65^, both known to induce mitochondrial gene expression, exhibited increased activity in hypoxia and were components of this transcription factor network. As such, targeted investigations using *in vitro* contraction-based models^66^ will be essential to elucidate the specific roles of these identified transcription factors in the hypoxia-induced transcriptional response to exercise.

Although HIF-1α was predicted to be activated by exercise even under normoxic conditions (**Fig. 5b**), the additional activity of its negative regulators (i.e., AHR and TP73) within the hypoxia-induced transcriptional network may counterbalance typical hypoxic responses, such as a shift toward greater reliance on anaerobic metabolism^13,67^, which are generally considered maladaptive adaptations to cardiorespiratory exercise in human skeletal muscle. This response could also contribute to acclimatisation^68^, potentially blunting the negative impact of future hypoxic exposures, including those encountered during exercise in normoxia. However, whether attenuation of hypoxic signalling is ultimately beneficial remains unclear. Notably, dampening of hypoxic signalling has been identified as an early adaptive process during repeated exercise sessions, resulting in reduced exercise-induced gene expression^59^. Taken together, the results of the present study do not allow a definitive conclusion regarding whether exercise in hypoxia adversely affects mitochondrial biogenesis. Future studies using similar experimental protocols but focusing on changes in mitochondrial protein abundance, volume density, and respiratory function following training are needed to clarify the functional impact of these gene expression differences between hypoxic exercise and intensity-matched exercise in normoxia.

While this study provides valuable insights into the transcriptomic response to exercise in both normoxia and hypoxia, several limitations should be acknowledged. Given the intensity-^28,69^ and volume-dependent^70^ nature of skeletal muscle adaptations, we employed HIIE to investigate the effects of hypoxia on exercise-induced gene expression. Whether the same genes would be differentially expressed, or whether hypoxia would similarly amplify their expression at +3 h and +24 h, may not generalise to other modalities, such as MICE or SIE. Furthermore, we recruited healthy, young males, and thus the findings may not be generalisable to females, older adults, or individuals with different health statuses or levels of cardiorespiratory fitness. The latter is particularly important, as higher fitness levels can attenuate exercise-induced gene expression^27^ and suppress HIF-1 signalling following training^59,71^. Participants also exhibited a range of V̇O_2peak_ values (30.4 to 54.9 mL·kg^-1^·min^-1^ in normoxia), meaning these results may not represent hypoxic responses in either sedentary or highly trained individuals^72^. Consequently, caution is warranted when interpreting these transcriptional changes, particularly when extrapolating them to predict future training-induced adaptations.

While exercise-induced changes in mRNA are often considered to forecast subsequent protein adaptations^73^, the evidence for a consistent predictive relationship is limited^27,74–76^. This is particularly relevant when comparing NA and NR, where mitochondrial gene expression and translation was upregulated at +24 h post-exercise in NA but not NR (**Fig. 3e**). This contrasts extensive evidence that higher exercise intensity induces a greater activation of mitochondrial biogenesis regulators^77^ and increased mitochondrial protein synthesis^78,79^ following a single session of exercise, and ultimately enhanced mitochondrial respiratory function following training^69^. Other explanations include the timing of muscle biopsies, as mounting evidence suggest peak gene expression for many mitochondrial-related genes varies considerably post-exercise^46^. Nonetheless, a recent comprehensive time-course study following HIIE reported minimal upregulation of most mitochondrial genes within 48 hours post-exercise, consistent with these findings^27^. As such, probing changes in gene expression may not be targeting the key regulatory step of mitochondrial adaptation, with post-transcriptional mechanisms potentially playing a larger role than commonly acknowledged^43,74–76^, further highlighting the need for additional studies using similar experimental protocols that focus on changes in mitochondrial characteristics following training.

In conclusion, our study demonstrates that normobaric hypoxia elicits a transcriptomic response to HIIE that is distinct from exercise matched by absolute and relative intensity in normoxia. In addition to amplifying a shared exercise response common to all conditions and an intensity-specific response observed only between hypoxic exercise and relative intensity-matched exercise in normoxia, hypoxia induced the differential expression of a substantially greater number of genes. The most pronounced differences were observed in mitochondrial-related pathways, which were downregulated at +24 h post-exercise under hypoxic, but not normoxic conditions. Overall, these findings suggest that exercise performed in normobaric hypoxia activates a unique transcriptional response compared with normoxia.

## Methods

### Participants and ethics approval

Ten healthy young males (28 ± 5 y; 175 ± 7 cm; 72.9 ± 10.0 kg; 23.9 ± 2.6 kg·m^-2^; 44.7 ± 8.2 mL·kg^-1^·min^-1^) volunteered for this study, as previously reported^11^. Participants underwent medical screening to exclude conditions that may have precluded their participation (e.g., cardiovascular, musculoskeletal, and/or metabolic disorders). After being informed of the study requirements, risks, and benefits, all provided written informed consent. The Victoria University Human Research Ethics Committee approved the study (HRE18-214), which conformed to the Declaration of Helsinki.

### Nutritional and physical activity controls

Participants were instructed to maintain their usual diet and physical activity levels throughout the study. They were advised to fast for two hours prior to any graded exercise test and completed diet and exercise questionnaires. Participants followed a controlled diet for the two days preceding and on the day of each HIIE muscle biopsy trial. Meal energy content was calculated using the Mifflin St-Jeor equation^80^, and nutritional composition of the packaged meals was verified using the FoodWorks (Xyris Pty Ltd, Australia) nutritional database to meet dietary requirements, consistent with previous methods^81^. Participants were requested to refrain from any strenuous exercise for 24 hours before exercise tests and 48 hours before HIIE muscle biopsy trial days.

### Graded exercise tests

Participants completed 4-6 graded exercise tests (GXTs) before the study to determine their peak power output (PPO), power at the lactate threshold (LT), and peak oxygen uptake (V̇O_2peak_) in both normoxia (FiO_2_ = 21%) and hypoxia (FiO_2_ = 14%). Participants performed a familiarisation GXT in both normoxia and hypoxia in a randomised order. They then completed a second GXT under both conditions, also randomised. If PPO differed by more than 10% between the initial two GXTs in either condition, a third GXT was conducted. The PPO and LT data from the two closest GXTs were then averaged to determine the exercise intensities for the HIIE sessions.

The GXT was performed on an electronically braked cycle ergometer (Lode Excalibur Sport) at 70-85 rpm cadence. Each test began at 20% of estimated PPO, which was calculated from demographic data, physical activity, height, and body mass data, as previously used in our lab^82^. Resistance increased by 10% of estimated PPO every 4 min, continuing until exhaustion, defined as the inability to maintain > 60 rpm, with a target duration of 40 min.

Blood samples were collected using single-use lancing needles (Accu-Chek Safe T-Pro Plus Lancet, Roche, Basel, Switzerland) and capillary tubes. Samples were taken at rest and immediately after each 4-min stage during 30-s breaks. Blood lactate concentration was determined using an automated analyser (YSI 2000 Glucose/Lactate Analyser, YSI, Ohio, USA). The LT was determined using the modified Dmax method (Dmax_MOD_) using Lactate-E software, incorporating power output, blood lactate, and heart rate data^83^. Expired gases were sampled every 15 s using a MOXUS Metabolic Cart System (AEI Technologies, Pennsylvania, USA) and V̇O_2peak_ was defined as the average of the four highest 15-s readings. Heart rate (Polar Electro Oy, Kempele, Finland) and a RPE using the Borg scale were recorded during the last 10 s of each stage.

### Exercise and muscle biopsy trials

Participants completed three separate exercise and muscle biopsy trials in a randomised order, each separated by one week: exercise in hypoxia (HY), exercise in normoxia matched to the absolute intensity of the session in hypoxia (NA), exercise in normoxia matched to the relative intensity of the session in hypoxia (NR).

Participants were asked to arrive at the laboratory in a fasted state between 7-8 am. An antecubital blood sample was collected before a *vastus lateralis* muscle biopsy (∼ 100 mg) was taken under local anaesthesia (5 mg·mL⁻¹ lidocaine) using the suction-modified Bergström technique (AgnTho’s, Sweden)^84^ by a qualified medical doctor. Excess blood, fat, and connective tissue were quickly removed from muscle biopsies before being frozen in liquid nitrogen and stored at -80 °C for RNA extraction. Additional antecubital blood samples and muscle biopsies were collected immediately after (+0 h), and at 3 hours (+3 h) and 24 hours (+24 h) post-exercise. For the exercise session in hypoxia, the PRE and +0 h blood samples and muscle biopsies were collected in hypoxic conditions. Muscle biopsies in subsequent trials were taken from alternating legs (e.g., left–right–left) to minimise sampling effects.

Each HIIE session comprised six 4-min intervals, interspersed with 2 min of passive recovery. In the HY session, the exercise work rate was set as the PPO achieved in hypoxia (PPO_H_) and the LT achieved in hypoxia (LT_H_), calculated as LT_H_ + 50% (PPO_H_ - LT_H_). In the NR session, the exercise work rate was based on PPO achieved in normoxia (PPO_N_) and LT achieved in normoxia (LT_N_), calculated as LT_N_ + 50% (PPO_N_ - LT_N_). For the NA session, the exercise work rate mirrored that of the HY session, equalling to LT_H_ + 50% (PPO_H_ - LT_H_).

Participants were permitted to engage in light activities (sitting or walking) and could drink water but were not allowed food for the 3 hours following the +0 h biopsy. Following the +3 h biopsy, standardised meals were provided before participants left the laboratory. They returned the next morning, fasted, for the +24 h biopsy.

### RNA-seq analysis

#### RNA extraction

Muscle samples (10-15 mg) were homogenised for 2 min at 30 Hz using the TissueLyser II (Qiagen, Germany), and total RNA was extracted utilising the Qiagen AllPrep DNA/RNA Kits (Qiagen, Germany), according to the manufacturer’s instructions.

RNA concentration and purity were assessed using a ND1000 Nanodrop spectrophotometer (Thermo Fisher Scientific, Wilmington, DE) and Agilent TapeStation 4150 (Agilent Technologies, USA). Samples with an RNA Integrity Number (RIN) ≥ 6 were deemed acceptable for further analysis.

#### RNA sequencing

Total RNA (1 μg) was processed using the NEBNext Ultra^TM^ RNA Library Prep Kit (Illumina NEB, USA), following the manufacturer’s protocols, with the following adjustments: 0.8× beads were used during the first purification step after second-strand synthesis; the adaptor was diluted at a 1:15 ratio; 0.7× beads were used for purification after adaptor ligand; 13 cycles of enrichment were performed; and a dual bead size selection (0.5× and 0.7×) was implemented for size selection of adaptor-ligated RNA. The samples were subsequently pooled and subjected to quality assessment through quantitative PCR (qPCR) and the Agilent 2100 Bioanalyzer (Agilent Technologies, USA).

The cDNA libraries were sequenced (100 bp, paired-end reads) using the Illumina NovaSeq 6000 (Illumina NEB, USA). Base sequence quality for all samples was > 91.49% bases above Q30. Transcriptome assembly was completed by alignment to the *Homo sapiens* genome (Build version HG38) using Hisat2 (v.2.0.4)^85^, with reads screened for the presence of any adaptor or over-represented sequences and cross-species contamination. Read counts mapped to known HGNC gene symbols were summarised, yielding 35-87 million clean reads per sample for downstream analysis.

#### Bioinformatic analysis

Count data was filtered to exclude lowly expressed genes, removing those with an average count per million (CPM) below 10 at any time point using the *edgeR* (v.4.4.1) package in R (v.4.5.2). The count data was then normalised for library size using the trimmed mean of M-values (TMM) method via the *edgeR* package in R^86^. Differential expression analysis was performed using the *edgeR* and *limma* (v.3.62.1) packages in R^87^. The voomLmfit function was applied to estimate mean-variance relationships, fit a model, and block for participant ID, while the eBayes function provided empirical Bayes moderation. Transcription factor activity was estimated using the *dorothea* (v.1.18.0) and *viper* (v.1.40.0) packages in R^33^. Differential transcription factor activity analysis was performed using *limma*, with lmFit used to fit a model and duplicateCorrelation to block for participant ID, while the eBayes function provided empirical Bayes moderation.

Upset plots were visualised using the *ComplexUpset* (v.1.3.3) and *ggplot2* (v.3.5.1) packages in R. Gene set enrichment analysis was completed using the *clusterProfiler* (v.4.14.6)^88^ package in R. Unsupervised hierarchical clustering with the ‘average’ method for Euclidean distances was used to visualise differentially expressed genes in a heatmap by the package *ComplexHeatmap* (v.2.22.0) in R. Transcription factor network protein-protein interactions were mapped using Gene MANIA (v.3.5.2)^36^ in Cytoscape (v.3.10.3)^89^. Additional plots were visualised using *ggplot2* and related packages in R.

#### Statistical analysis

Differentially expressed genes, gene ontology terms, or transcription factor activities were considered significant at an adjusted p-value < 0.05, using the Benjamini-Hochberg (BH) method. For the comparison of exercise prescriptions, a linear mixed-effects model was fitted using the *lme4* package (v1.1-37) in R, with a fixed effect for condition and random intercepts for participant ID. Post-hoc pairwise comparisons between groups were performed by estimated marginal means using the *emmeans* (v.1.10.6) package in R, with adjustment for multiple comparisons using the Tukey’s honest significance difference method. Degrees of freedom were estimated using the Kenward-Roger approximation. For the analysis of glucose and lactate, a linear mixed-effects model was fitted using the *lme4* package in R, with fixed effects for the interaction between time and sex, and random intercepts for participant ID. Post-hoc pairwise comparisons between groups were performed by estimated marginal means using the *emmeans* package in R, with adjustment for multiple comparisons using the Bonferroni-Holm method. Degrees of freedom were estimated using the Kenward-Roger approximation. Significance was defined as an adjusted p-value < 0.05 for all post-hoc pairwise comparisons. All values are reported or visualised as means ± SD, unless otherwise specified. Additional statistical details are provided in the figure legends.

## Acknowledgments

We wish to thank all the participants for their time and commitment in the present study. The authors would also like to thank the technical team at Victoria University for their assistance throughout the study. We also thank the creators of all the repositories, software, and R packages used in this study, whose contributions were invaluable but could not be individually cited due to reference limitations. This study was funded by an international collaborative grant from Beijing Sport University, China (2018GJ005) to Y.L. and X.Y., a Key Science and Technology Project grant from Lhasa, China (LSKJ202634) to J.L. and N.O., and an Australian Research Council (ARC) Discovery Project grant (DP200103542) awarded to D.J.B.

## Author contributions

D.J.B., Y.L, X.Y., J.L., and D.F.T. conceptualised the study and devised the study methodology. J.L., J.K., Z.W., N.Z., M.M.A., and X.Y. delivered the exercise sessions and performed sample collection. J.L., J.K., K.C., A.G., Y.L., and X.Y. performed the sample processing. RNA-seq analysis was performed at Biomarker Technologies (BMKGENE), China. D.F.T. performed statistical and bioinformatic analysis. D.F.T. delivered the visualisation. J.L., D.F.T., Y.L., D.J.B., and X.Y. wrote the initial draft of the manuscript. J.L., D.F.T., Y.L., D.J.B., and X.Y. have primary responsibility for final content. Data collection took place at Victoria University. Muscle analysis took place at BMKGENE and Victoria University. All persons designated as authors qualify for authorship, and all those qualifying for authorship are listed. All authors have read and approved the final manuscript.

---

**Supplementary Fig. 1.**
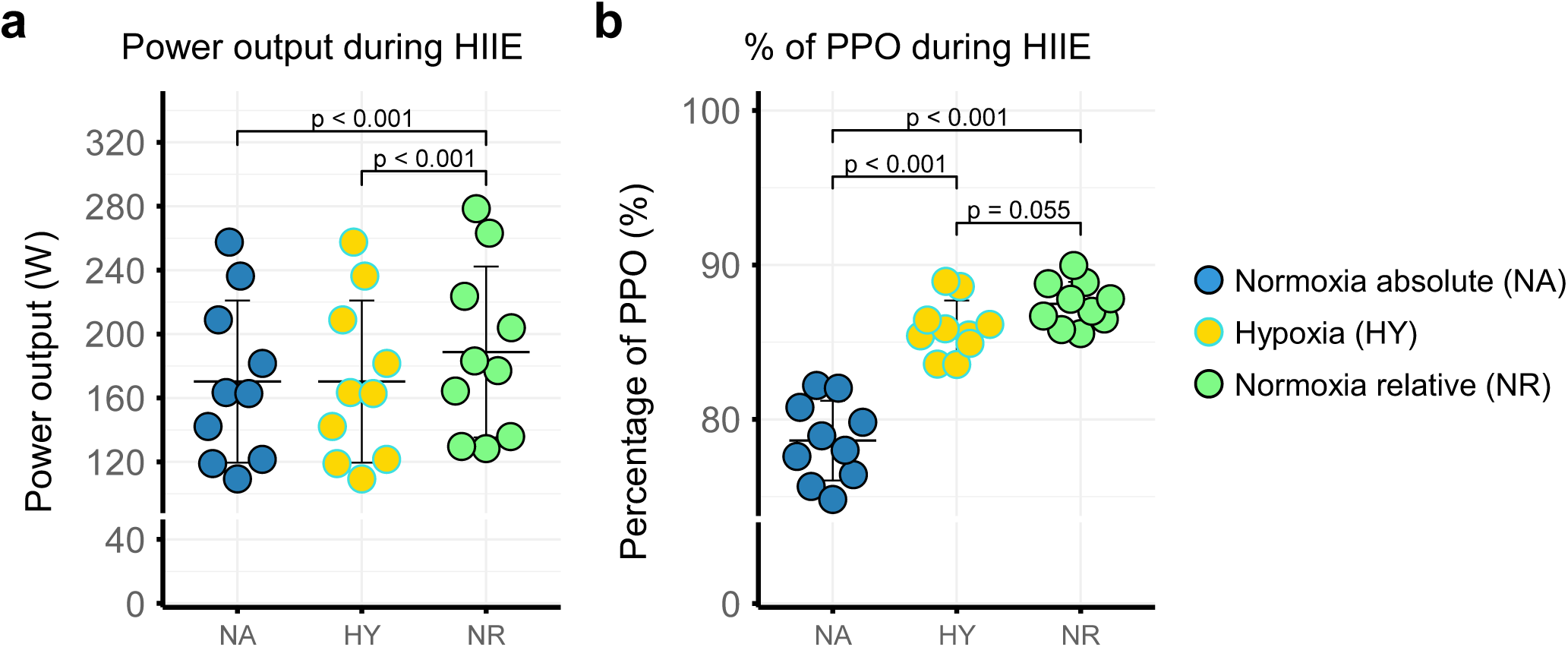
Differences in exercise prescription between NA, HY, and NR conditions. **a** Differences in power output between normoxia absolute (NA), hypoxia (HY), and normoxia relative (NR). **b** Differences in the percentage of peak power output (PPO) between NA, NR, and HY within normoxia and hypoxia (depending on the respective condition).

**Supplementary Fig. 2.**
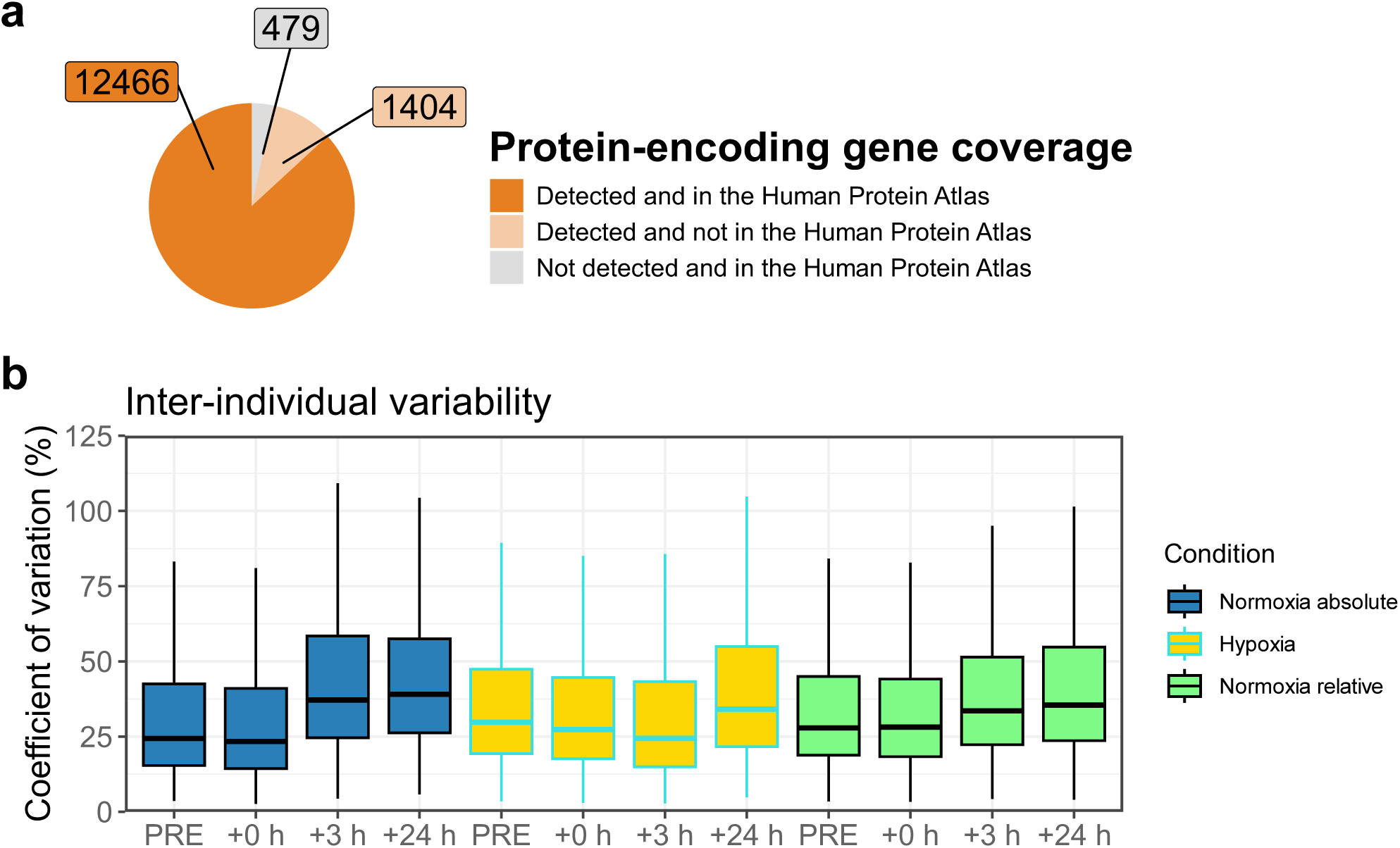
Protein-coding coverage, and inter- and intra-individual differences. **a** Protein-coding transcript depth compared to the skeletal muscle database of the Human Protein Atlas^26^. **b** Inter-individual variability of all detected genes using count per million (CPM) values at each time point. Each box represents the interquartile range (IQR) and median of the coefficient of variation. Each whisker representing 1.5 * IQR for both directions. Outliers are not displayed.

**Supplementary Fig. 3.**
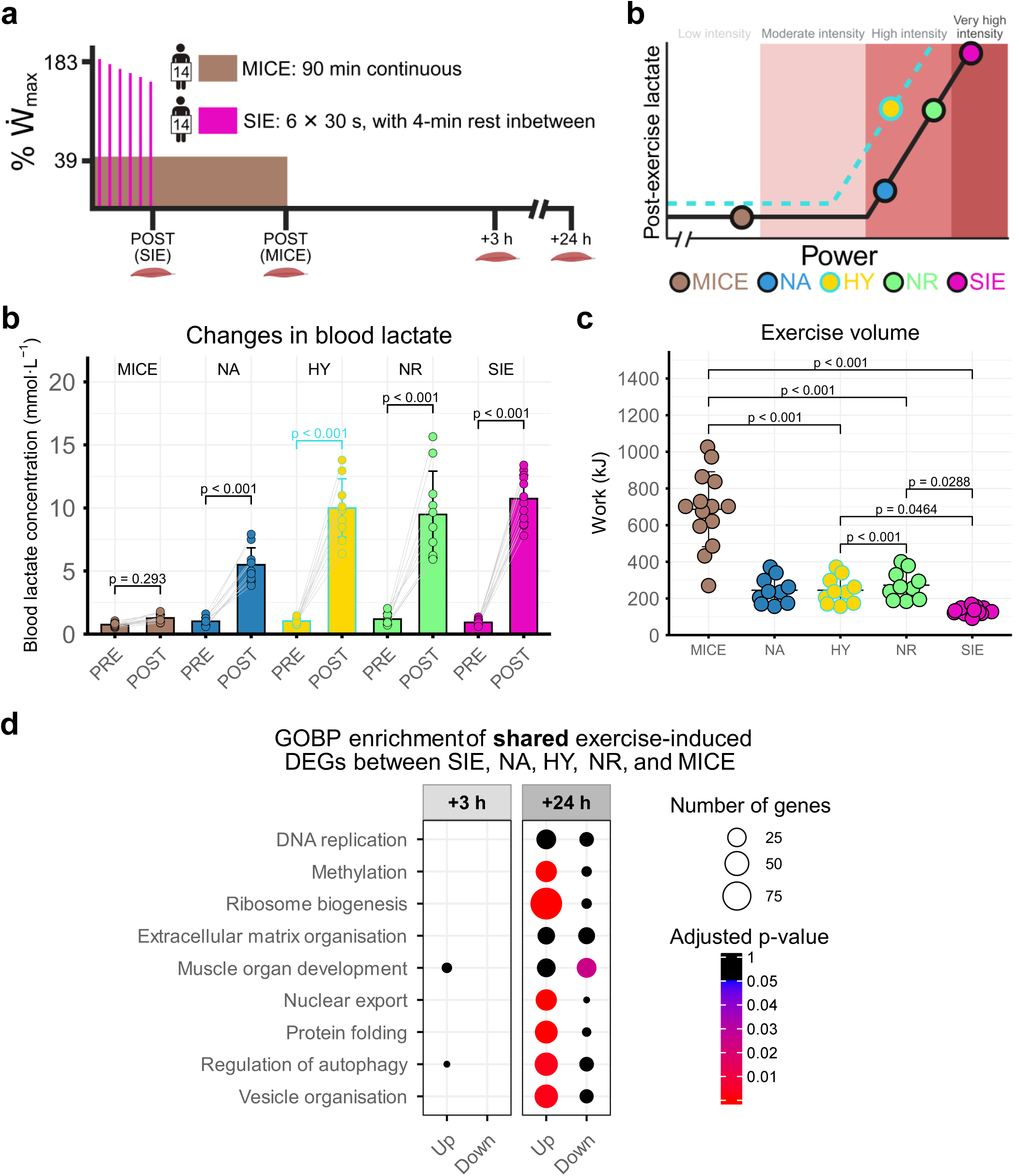
Shared differentially expressed genes across NA, HY, and NR and their overlap with moderate- and sprint-interval exercise. **a** Exercise prescriptions used in Botella *et al*.^31^ for moderate-intensity continuous exercise (MICE) and sprint-interval exercise (SIE). Ẇ_max_: maximum power output. **b** Changes in antecubital blood lactate following HIIE in NA, HY, and NR, as well as SIE and MICE in Botella *et al*. **c** Differences in exercise volume, calculated in kJ, between exercise groups in this study and SIE and MICE in Botella *et al*. **d** Upset plot showing upregulated and downregulated differentially expressed genes (DEGs) at each time point relative to PRE across all conditions (NA, HY, and NR) and MICE and SIE from Botella *et al*., with only genes quantified in both studies included in the comparison. The proportion of each set that is protein-coding is indicated. Minimum set size displayed is 1. ncRNA: non-coding RNA.

**Supplementary Fig. 4.**
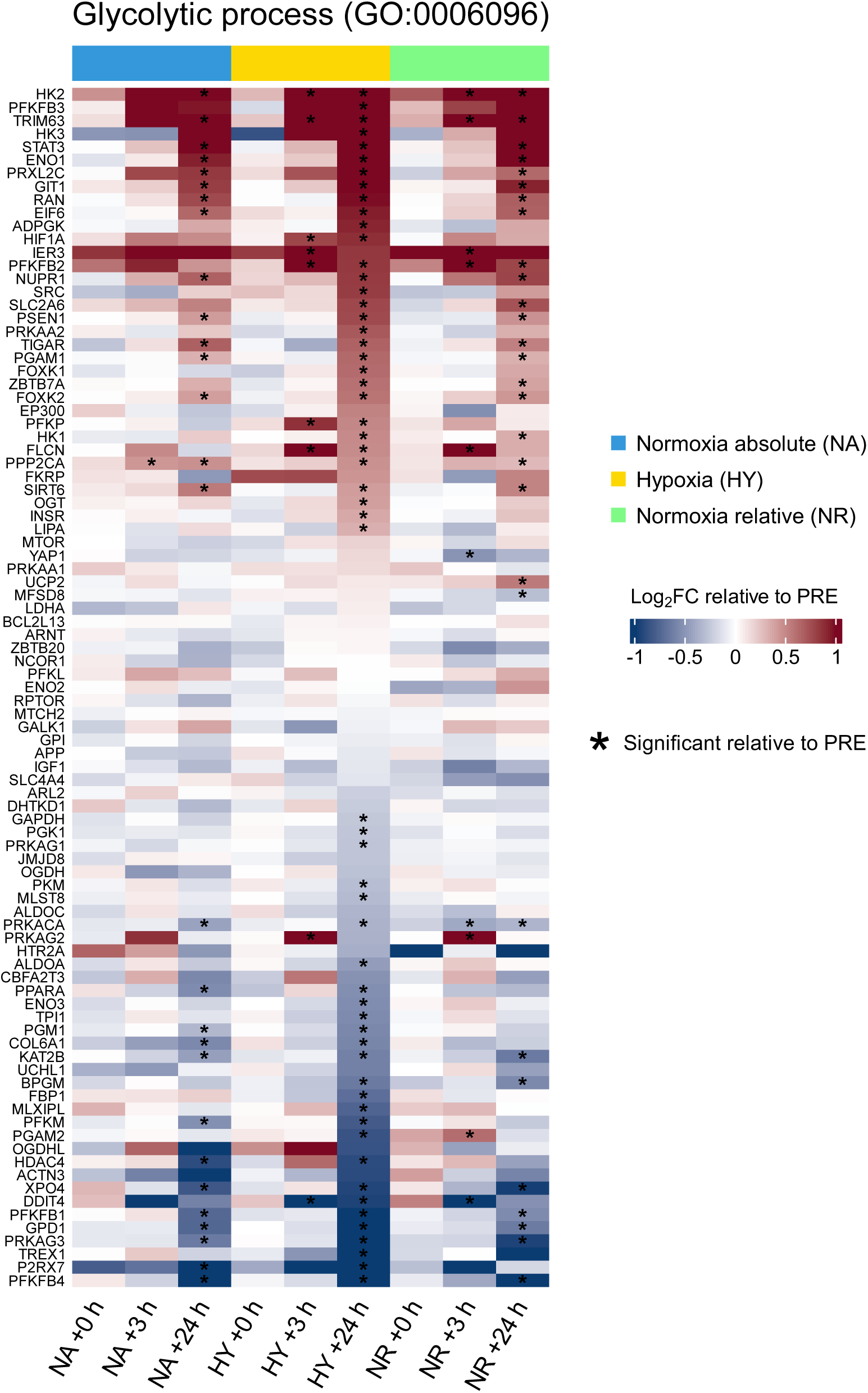
Hypoxic-specific regulation of genes involved in glycolytic processes. Heatmap of genes quantified in the gene ontology biological process ‘glycolytic process’ (GO:0006096). Significance was determined with an adjusted p-value < 0.05 (Benjamini-Hochberg) and is indicated by * for each combination of condition and time point.

